# Topology, cross-frequency, and same-frequency band interactions shape the generation of phase-amplitude coupling in a neural mass model of a cortical column

**DOI:** 10.1101/023291

**Authors:** Roberto C. Sotero

## Abstract

Phase-amplitude coupling (PAC), a type of cross-frequency coupling (CFC) where the phase of a low-frequency rhythm modulates the amplitude of a higher frequency, is becoming an important indicator of information transmission in the brain. However, the neurobiological mechanisms underlying its generation remain undetermined. A realistic, yet tractable computational model of the phenomenon is thus needed. Here we propose a neural mass model of a cortical column, comprising fourteen neuronal populations distributed across four layers (L2/3, L4, L5 and L6). The conditional transfer entropies (cTE) from the phases to the amplitudes of the generated oscillations are estimated by means of the conditional mutual information. This approach provides information regarding directionality by distinguishing PAC from APC (amplitude-phase coupling), i.e. the information transfer from amplitudes to phases, and can be used to estimate other types of CFC such as amplitude-amplitude coupling (AAC) and phase-phase coupling (PPC). While experiments often only focus on one or two PAC combinations (e.g., theta-gamma or alpha-gamma), we found that a cortical column can simultaneously generate almost all possible PAC combinations, depending on connectivity parameters, time constants, and external inputs. We found that the strength of PAC between two populations was strongly correlated with the strength of the effective connections between them and, on average, did not depend upon the presence or absence of a direct (anatomical) connection. When considering a cortical column circuit as a complex network, we found that neuronal populations making indirect PAC connections had, on average, higher local clustering coefficient, efficiency, and betweenness centrality than populations making direct connections and populations not involved in PAC connections. This suggests that their interactions were more efficient when transmitting information. Since more than 60% of the obtained interactions represented indirect connections, our results highlight the importance of the topology of cortical circuits for the generation on of the PAC phenomenon. Finally, our results demonstrated that indirect PAC interactions can be explained by a cascade of direct CFC and same-frequency band interactions, suggesting that PAC analysis of experimental data should be accompanied by the estimation of other types of frequency interactions for an integrative understanding of the phenomenon.

## 1. Introduction

It has been hypothesized that phase-amplitude coupling (PAC) of neurophysiological signals plays a role in the shaping of local neuronal oscillations and in the communication between cortical areas (Canolty and Knight 2010). PAC occurs when the phase of a low frequency oscillation modulates the amplitude of a higher frequency oscillation. A classic example of this phenomenon was demonstrated in the CA1 region of the hippocampus (Bragin, Jando et al. 1995), where the phase of the theta band modulated the power of the gamma-band. Computational models of the theta-gamma PAC generation in the hippocampus have been proposed (Kopell, Boergers et al. 2010) and are based on two main types of models. The first type of models consists of a network of inhibitory neurons (I-I model) (White, Banks et al. 2000), whereas the second model is based on the reciprocal connections between networks of excitatory pyramidal cells and inhibitory neurons (E-I model) (Tort, Rotstein et al. 2007, Kopell, Boergers et al. 2010). In such models, fast excitation and delayed feedback inhibition alternate, and with appropriate strength of excitation and inhibition, oscillatory behavior occurs. When the gamma activity produced by the E-I or I-I models is periodically modulated by a theta rhythm imposed by either an external source or theta resonant cells within the network (White, Banks et al. 2000), a theta-gamma PAC is produced. Recently, the generation of theta-gamma PAC was studied (Onslow, Jones et al. 2014) using a neural mass model (NMM) proposed by Wilson and Cowan (Wilson and Cowan 1972). In NMMs, spatially averaged magnitudes are assumed to characterize the collective behavior of populations of neurons of a given type instead of modeling single cells and their interactions in a realistic network (Wilson and Cowan 1972, Jansen and Rit 1995). Specifically, the Wilson and Cowan model consists of excitatory and inhibitory neural populations which are mutually connected.

While the models mentioned above have improved our understanding of the physiological mechanism that gives rise to theta-gamma PAC, we lack modeling insights into the generation of PAC involving other frequency pairs (Sotero 2015). This is critical because experimental studies have shown that the PAC phenomenon is not restricted to either the hippocampus or to theta-gamma interactions. In fact, PAC has been detected in pairs involving all possible combinations of low and high frequencies: delta-theta (Lakatos, Shah et al. 2005), delta-alpha (Cohen, Elger et al. 2009, Ito, Maldonado et al. 2013), delta-beta (Cohen, Elger et al. 2009, Nakatani, Raffone et al. 2014), delta-gamma (Gross, Hoogenboom et al. 2013, Lee and Jeong 2013, Nakatani, Raffone et al. 2014, Szczepanski, Crone et al. 2014, Florin and Baillet 2015), theta-alpha (Cohen, Elger et al. 2009), theta-beta (Cohen, Elger et al. 2009, Nakatani, Raffone et al. 2014), theta-gamma (Lakatos, Shah et al. 2005, Demiralp, Bayraktaroglu et al. 2007, Wang, Li et al. 2011, Durschmid, Zaehle et al. 2013, Lee and Jeong 2013, McGinn and Valiante 2014, Florin and Baillet 2015), alpha-beta (Sotero, Bortel et al. 2015), alpha-gamma (Osipova, Hermes et al. 2008, Voytek, Canolty et al. 2010, Spaak, Bonnefond et al. 2012, Wang, Saalmann et al. 2012), and beta-gamma (Wang, Saalmann et al. 2012, de Hemptinne, Ryapolova-Webb et al. 2013). Furthermore, although experimental studies usually focus on one or two PAC combinations, most of the combinations mentioned above can be detected in a single experiment (Sotero, Bortel et al. 2015). This suggests a diversity and complexity of the PAC phenomenon that has not been incorporated into current computational models. Similarly, there is a need for further improvement in the mathematical methods used to detect PAC. Although a large number of methods have been proposed (Penny, Duzel et al. 2008, Tort, Komorowski et al. 2010), no gold standard has emerged.

In this work, we propose a neural mass model of a cortical column that comprises 4 cortical layers and 14 neuronal populations and study the simultaneous generation of all PAC combinations mentioned above. To estimate PAC we use a measure of the information transfer from the phase of the low frequency rhythm to the amplitude of the higher frequency oscillation, which is known as conditional transfer entropy (cTE) (Lizier, Heinzle et al. 2011). This multivariate approach provides information about the directionality of the interactions, thus distinguishing PAC from the information transfer from the amplitude to the phases (i.e. amplitude-phase coupling, or APC) which has been experimentally detected (Jiang, Bahramisharif et al. 2015). This done in contrast to previous methods which were either based on pairwise correlations between the selected phase and amplitude (Canolty, Edwards et al. 2006, Penny, Duzel et al. 2008), or provided directionality using pairwise approaches (Jiang, Bahramisharif et al. 2015), or were multivariate but did not provide directionality (Canolty, Cadieu et al. 2012). By estimating cTE from phases to amplitudes, we obtain a clearer view of the mechanisms underlying the generation of PAC in the cortical column which allows us to study the link between anatomical and effective PAC structure. In the examples shown in this paper, the neuronal populations modeled have natural frequencies in the theta, alpha and gamma bands. However, due to the effective connectivity between populations, oscillations in the delta and beta bands appear and result in PAC involving these frequencies. We focused on three combinations (delta-gamma, theta-gamma, and alpha-gamma) and explored how changes in model parameters such as the strength of the connections, time constants or external inputs strengthen or weaken the PAC phenomenon. We found that more than 60% of the obtained PAC interactions result from indirect connections and that, on average, these interactions have the same strength as direct (anatomical) connections. The cortical column circuit was analyzed as a complex network and three different local topological measures were computed: the clustering coefficient (*C_m_*), the efficiency (*E_m_*) and betweenness centrality (*B_m_*) which quantify how efficiently the information is transmitted within the network. According to our results, neuronal populations sending direct PAC connections had higher local *C_m_*, *E_m_*, and *B_m_* coefficients, than populations receiving the PAC connection and populations not involved in PAC interactions. This suggests that the topology of cortical circuits plays a central role in the generation of the PAC phenomenon.

Finally, although this paper focuses on the PAC phenomenon, in order to study the generation of indirect PAC connections we also estimated other types of cross-frequency coupling such as APC, amplitude-amplitude coupling (AAC), and phase-phase coupling (PPC), as well as interactions within the same frequency band (or same-frequency coupling, SFC), and used these as predictors of indirect PAC in a linear regression analysis. We demonstrated that indirect PAC connections can be predicted by a cascade of direct CFC and SFC interactions, suggesting that PAC analysis of experimental data should be accompanied by the estimation of other types of interactions for an integrative understanding of the phenomenon.

A list of the abbreviations used in this paper is presented in Table 1.

**Table 1.** List of abbreviations.

| Abbreviation | Meaning |
| --- | --- |
| AAC | Amplitude-amplitude coupling |
| APC | Amplitude-phase coupling |
| CFC | Cross-frequency coupling |
| cMI | Conditional mutual information |
| cTE | Conditional transfer entropy |
| ECoG | Electrocorticography |
| EEG | Electroencephalography |
| ESC | Envelope-to-signal correlation |
| FS | Fast-spiking |
| IB | Intrinsically bursting |
| LFP | Local field potential |
| LTS | Low-threshold |
| Midx | Modulation index |
| NMM | Neural mass model |
| PAC | Phase-amplitude coupling |
| PFC | Phase-frequency coupling |
| PPC | Phase-phase coupling |
| PSP | Postsynaptic potential |
| RS | Regular spiking |
| SFC | Same-frequency coupling |

## 2. Methods

### 2.1. A neural mass model of a cortical column

Figure 1 shows the proposed model obtained by distributing four cell classes in four cortical layers (L2/3, L4, L5, and L6). This produced 14 different neuronal populations, since not all cell types are present in each layer (Neymotin, Jacobs et al. 2011). Excitatory neurons were either regular spiking (RS) or intrinsically bursting (IB), and inhibitory neurons were either fast-spiking (FS), or low-threshold spiking (LTS) neurons. The evolution of each population dynamics rests on two mathematical operations. Post-synaptic potentials (PSP) at the axonal hillock were converted into an average firing rate using the sigmoid function:

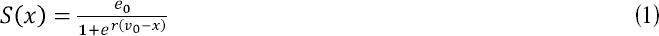
where the variable *x* represents the PSP and parameters *e_0_, v*_0_ and *r* represent the maximal firing rate, the PSP corresponding to the maximal firing rate e_0_, and the steepness of the sigmoid function, respectively. The second operation was the conversion of firing rate at the soma and dendrites into PSP, which was done by means of a linear convolution with an impulse response *g(t)* given by:

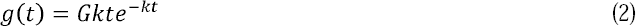

where *G* controls the maximum amplitude of PSP and *k* is the sum of the reciprocal of the average time constant (Jansen and Rit 1995). The convolution model with impulse response (2) can be transformed into a second order differential equation (Jansen and Rit 1995, Sotero, Trujillo-Barreto et al. 2007). The temporal dynamics of the average PSP in each neuronal population *x_m_* can then be obtained by solving a system of 14 second order differential equations:

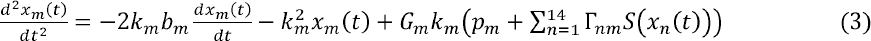

where *n* = 1,…,14 and *m* = 1,…,14. The populations are numbered from 1 to 14 following the order: [L2RS, L2IB, L2LTS, L2FS, L4RS, L4LTS, L4FS, L5RS, L5IB, L5LTS, L5FS, L6RS, L6LTS, L6FS]. Notice that layer 2/3 was simply labelled as L2. As can be seen in (3), neuronal populations interact via the connectivity matrix Γ*_nm_*. This is an ‘anatomically constrained’ effective connectivity matrix (Sotero, Bortel et al. 2010) in the sense that its elements represent anatomical (i.e., direct) connections, but their strength (except the ones set to zero) can vary with a condition or task. Inputs from neighboring columns are accounted for via *p_m_*, which can be any arbitrary function, including white noise (Jansen and Rit 1995). Thus, equation (3) represents a system of 14 random differential equations (Carbonell, Jimenez et al. 2005).

**Figure 1.**
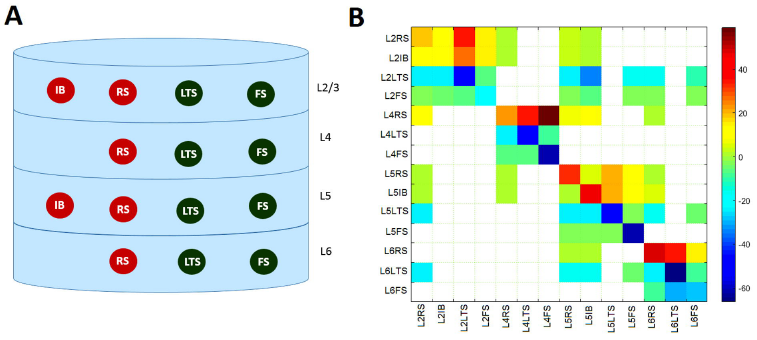
Proposed neural mass model of the cortical column. A) Layer distribution of the four neuronal types. The excitatory populations are the intrinsically bursting (IB), and the regulatory spiking (RS). The inhibitory populations are low-threshold spiking (LTS) and fast spiking (FS). B) Connectivity matrix values used for coupling the 14 populations modeled. Negative values correspond to inhibitory connections.

The ‘damping’ parameter *b_m_* critically determines the behavior of the system. If the connections between the populations are set to zero (Γ*_nm_* — 0,*n* ≠ *m*), then for *b_m_* > 1 (overdamped oscillator) and *b_m_* = 1 (critically damped oscillator), each neuronal population will evolve to a fixed point 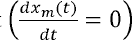 without oscillating. If *b_m_* < 1 (underdamped oscillator), each population is capable of producing oscillations even if the inter-population coupling is set to zero. The case *b_m_* = 1 corresponds to the Jansen and Rit model (Jansen and Rit 1995), which has been extensively used in the literature (David and Friston 2003, Grimbert and Faugeras 2006, Zavaglia, Astolfi et al. 2006, Sotero, Trujillo-Barreto et al. 2007, Sotero and Trujillo-Barreto 2008, Valdes-Sosa, Sanchez-Bornot et al. 2009, Ursino, Cona et al. 2010, Zavaglia, Cona et al. 2010). Thus, in this model, an individual population is not capable of oscillating, and the balance between excitation and inhibition is what produces oscillatory behavior that mimics observed Electroencephalography (EEG) signals. It should be noted that realistic models of a single inhibitory neural population are able to produce oscillations (Wang and Buzsaki 1996), but that excitatory populations were believed to only produce unstructured population bursts (Buzsáki 2006). This view has been challenged recently by both experimental and computational studies (Allene, Cattani et al. 2008, Tattini, Olmi et al. 2012). To account for the possibility of oscillatory activity in single populations, we introduced the parameter *b_m_* with values *b_m_* < 1. Tables 1 and 2 present the parameters of the model and their interpretation. As shown in table 2, FS populations have the fastest time constants, followed by IB, RS, and LTS, in that order.

**Table 2.**
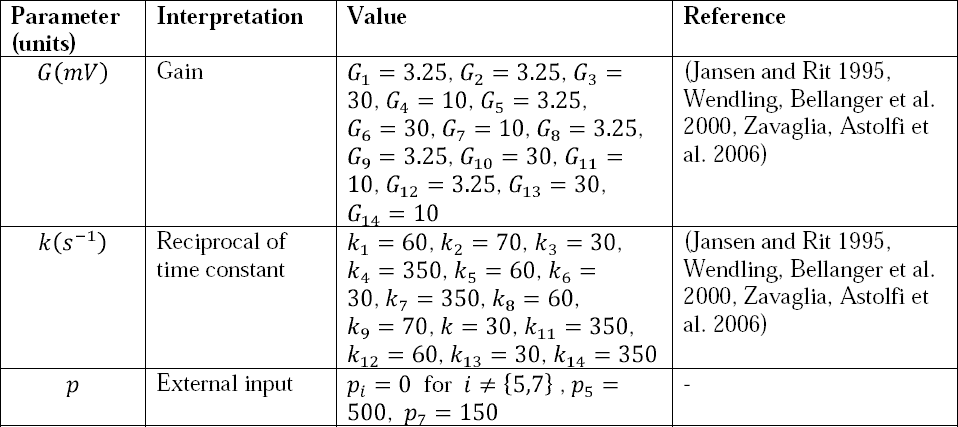
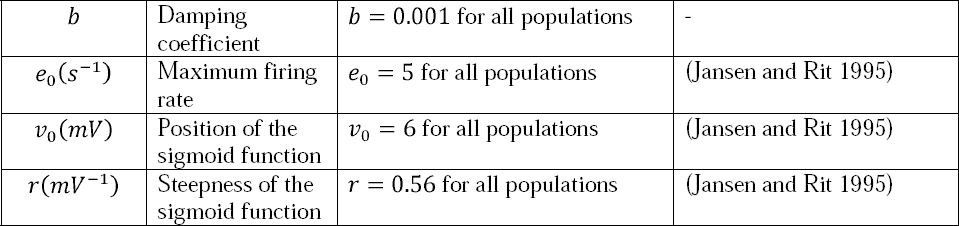
Values and physiological interpretations of model parameters for the 14 neuronal populations.

| Parameter (units) | Interpretation | Value | Reference |
| --- | --- | --- | --- |
| $G(mV)$ | Gain | $G_1 = 3.25, G_2 = 3.25, G_3 = 30, G_4 = 10, G_5 = 3.25, G_6 = 30, G_7 = 10, G_8 = 3.25, G_9 = 3.25, G_{10} = 30, G_{11} = 10, G_{12} = 3.25, G_{13} = 30, G_{14} = 10$ | (Jansen and Rit 1995, Wendling, Bellanger et al. 2000, Zavaglia, Astolfi et al. 2006) |
| $k(s^{-1})$ | Reciprocal of time constant | $k_1 = 60, k_2 = 70, k_3 = 30, k_4 = 350, k_5 = 60, k_6 = 30, k_7 = 350, k_8 = 60, k_9 = 70, k_{10} = 30, k_{11} = 350, k_{12} = 60, k_{13} = 30, k_{14} = 350$ | (Jansen and Rit 1995, Wendling, Bellanger et al. 2000, Zavaglia, Astolfi et al. 2006) |
| $p$ | External input | $p_i = 0$ for $i \neq \{5,7\}, p_5 = 500, p_7 = 150$ | - |
| $b$ | Damping coefficient | $b = 0.001$ for all populations | - |
| $e_0(s^{-1})$ | Maximum firing rate | $e_0 = 5$ for all populations | (Jansen and Rit 1995) |
| $v_0(mV)$ | Position of the sigmoid function | $v_0 = 6$ for all populations | (Jansen and Rit 1995) |
| $r(mV^{-1})$ | Steepness of the sigmoid function | $r = 0.56$ for all populations | (Jansen and Rit 1995) |

### 2.2 Estimation of phase-amplitude coupling

Several mathematical methods for detecting PAC have been proposed (Penny, Duzel et al. 2008, Canolty and Knight 2010, Tort, Komorowski et al. 2010, Canolty, Cadieu et al. 2012, Jiang, Bahramisharif et al. 2015), although each yields advantages and caveats, such that no gold standard for the detection of PAC has emerged. Although diverse, the basis for these methods is to test the correlation between the instantaneous phase of a lower frequency rhythm and the instantaneous amplitude of the higher frequency rhythm. To compute any one of these measures, signals generated with the model (3) need to be band-pass filtered into different frequency bands. In this paper we use the following bands: delta (0.1−4 Hz), theta (4−8 Hz), alpha (8−12 Hz), beta (12−30 Hz), and gamma (30−120 Hz). To this end, we designed FIR filters using MATLAB’s signal processing toolbox function *firls.m*. To remove any phase distortion, the filters were applied to the original time series in the forward and then the reverse direction using MATLAB’s function *filtfilt.m* (Penny, Duzel et al. 2008). The analytic representation *y_mk_(t)* of each filtered signal *x_mk_* (where *m* = 1,..,5 stands for the index of the frequency band, and *k* = 1,..,14, indexes the neuronal populations) was obtained using the Hilbert transform Hilbert ( x_m/c_(t)):

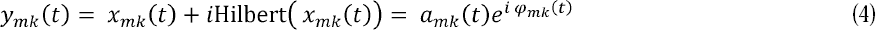

where *a_mk_(t)* and *φ_mk_(t)* are the instantaneous amplitudes and phases, and *i* is the imaginary number. Amplitudes were normalized by subtracting the temporal mean and dividing the result by the temporal standard deviation to create the set of normalized band-passed signals. Normalization was done to facilitate comparison between different frequency bands.

Two examples of PAC measures frequently used in the literature are the modulation index (Midx) (Canolty, Edwards et al. 2006) and the envelope-to-signal correlation (ESC) (Penny, Duzel et al. 2008):

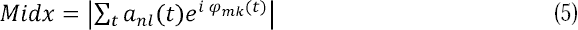

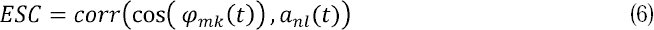
where subindexes *m* and *n* corresponds to different frequency bands and subindexes *k* and *l* correspond to different neuronal populations. However, ESC and Midx are pairwise measures of the correlation between phases and amplitudes and thus cannot detect directionality in the interaction. Measures such as cTE (Lizier, Heinzle et al. 2011) which are based on the information transmitted between signals should provide a clearer picture of the mechanisms generating PAC than correlation-based measures. cTE can be computed using the conditional mutual information (cMI) measure (MacKay 2003). First, we define the cMI between the phase *φ_mk_* and the amplitude *a_nl_*, given all the other phases (Φ) and amplitudes (A) as:

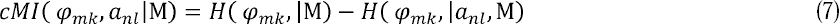

where *M* = [Φ, A] is a matrix comprising all phases and amplitudes in all populations, except *φ_mk_* and *a_nl_* and the two terms at the right side of the equation are conditional entropies (MacKay 2003). To compute cMI we use a toolbox (http://www.cs.man.ac.uk/˜pococka4/MIToolbox.html) which computes several information measures using the conditional likelihood maximization algorithm (Brown, Pocock et al. 2012). cMI does not provide information about the directionality of the coupling between phases and amplitudes, which is a problem because both theoretical (Daffertshofer and van Wijk 2011) and experimental (Jiang, Bahramisharif et al. 2015) studies indicate the possibility of an information flow from amplitudes to phases. On the other hand, cTE provides directionality by estimating the cMI between one signal (the phase in our case) and the other signal (the amplitude) shifted *δ* steps into the future. In this paper, to estimate *cTE* from the phase to the amplitude (denoted as *cTE_φ_mk_→a_nl__*), we compute *cMI* for *N* different *δ*s and average the results (Palus, Komarek et al. 2001, Palus and Stefanovska 2003, Lizier, Heinzle et al. 2011):

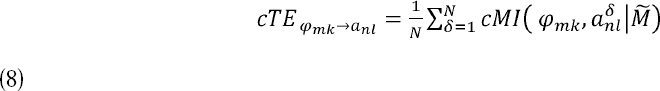

where 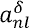 is derived from the amplitude time series *a_nl_* at *δ* steps into the future, i.e. 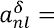 *a_nl_*(*t* + *δ*), and 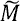 is a matrix comprising all phases and amplitudes in all populations, except (*φ_mk_*. In this paper we use *N* = 100. Since we use a time step of 10^-4^ s in all simulations, we are averaging the *cMI* up to a period of 10 ms into the future.

A significance value can be attached to any of the above measures by means of a surrogate data approach (Canolty, Edwards et al. 2006, Penny, Duzel et al. 2008), where we offset *φ_mk_* and *a_nl_* by a random time lag. We can thus compute 1000 surrogate Midx, ESC, cMI and cTE values. From the surrogate dataset we first compute the mean μ and standard deviation *σ*, and then compute a z-score as:

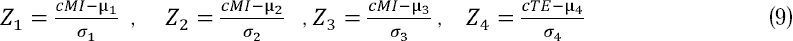

The p-value that corresponds to the standard Gaussian variate is also computed. Z values satisfying |*Z*| > 1.96 are significant with *α* = 0.05. Masks of zeros (for non-significant Z values) and ones (for significant Z-values) are created and multiplied to Midx, ESC, cMI, and cTE. Finally, a multiple comparison analysis based on the False Discovery Rate (Storey and Tibshirani 2003) is performed using the computed p-values.

### 2.3 Nonlinear correlation coefficient

Given the nonlinear nature of the PAC phenomenon, studying the link between the parameters of the model and the strength of PAC cannot be done with the Pearson correlation coefficient, which measures the linear correlation between two variables and is therefore not appropriate. Nonlinear measures are thus required. The underlying idea is that if the value of the variable X is considered as a nonlinear function of the variable Y, the value of Y given X can be predicted according to a nonlinear regression (Pereda, Quiroga et al. 2005). In this paper, we computed the nonlinear regression by fitting *y(t)* with a Fourier series:

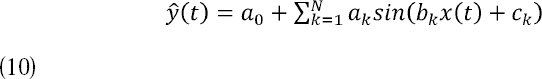

where we used *N* = 10. The nonlinear correlation coefficient *r_nl_* is then the value of the linear correlation between *y(t)* and the predicted signal *ŷ(t)*.

### 2.4. Topological properties of the cortical column network

Complex network analysis have proven useful for studying the relationship between structure and function in brain networks (Honey, Kotter et al. 2007). In this paper we are interested in studying how the topology of the connectivity matrix Γ*_nm_* influences the PAC phenomenon. Specifically, we want to answer the question of whether the populations involved in direct and indirect PAC interactions present the same topological properties. This means we need to focus on local properties of the network instead of global ones. In this paper we are going to compute three such properties: the local clustering coefficient, the local efficiency, and the local betweenness centrality, for the sending and receiving nodes involved in each direct or indirect PAC interaction.

In this section we are not going to distinguish between inhibitory and excitatory connections, and the analysis will be done to the absolute value of the connectivity matrix: *W* = |Γ*_nm_*|. Nodes of a network can be characterized by the structure of their local neighborhood. The concept of clustering of a network refers to the tendency to form cliques in the neighborhood of any given node (Watts and Strogatz 1998). This means that if node *m* is connected to node *n*, while at the same time node *n* is connected to node *s*, there is a high probability that node *m* is also connected to node *s*.

Let *A* = {*a_mn_*} be the directed adjacency matrix (Albert and Barabasi 2002) of the network (*a^mn^* = 1 when there is a connection from *m* to *n, a_mn_* = 0 otherwise). Let also 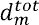 be the total degree of node *m*, and 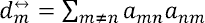. The local clustering coefficient of node *m* for weighted networks is (Fagiolo 2007):

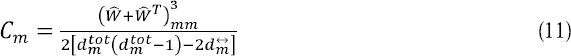

where 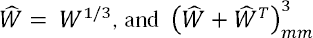 is the *m*th element of the main diagonal of 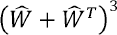.

The second measure we are going to compute is the local efficiency, calculated as (Latora and Marchiori 2001, Rubinov and Sporns 2010):

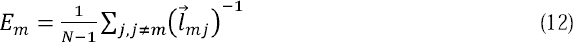

where 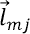 is the shortest weighted path length from *m* to *j*. Thus, *E_m_* is inversely related to the path length, and measures how efficiently the network exchanges information on a local scale.

To account quantitatively for the role of nodes that can be crucial for connecting different regions of the network by acting as bridges, the concept of betweenness centrality was introduced (Newman, Barabási et al. 2006). The local weighted betweenness centrality of node *m* is computed as (Rubinov and Sporns 2010):

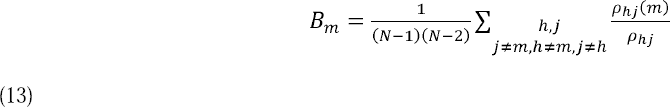

where *ρ_hj_* is the number of shortest paths between *h* and *j* that pass through *m*. A node with high centrality is thus crucial to efficient communication.

To compute the *C_m_*, *E_m_*, and *B_m_* measures we use Matlab functions provided in the brain connectivity toolbox (https://sites.google.com/site/bctnet/).

## 3. Results

### 3.1 Detecting PAC: control analysis

We connected three excitatory neuronal populations, labeled 1, 2 and 3 (Figure 2A and 2B). The temporal dynamics of the three populations are described by a system of random differential equations identical to (3), but with n=1:3 and m=1:3. As shown in Figure 2A, there is no connection between populations 1 and 3 and both are driven by population 2. The parameters used in this simulation were: *G*_1_ = 3.25, *G*_2_ = 4, *G*_2_ = 3.25, *G*_3_ = 4, *k*_1_ = 330, *k*_2_ = 30, *k*_3_ = 400. Inputs *p*_1_, *p*_2_, and *p*_3_ were white noise processes with mean 0 and standard deviations: *σ*_1_ = *σ*_2_ = *σ*_3_ = 3. Simulated data were generated by numerically integrating system (3). To do this, the local linearization method for random systems was used (Carbonell, Jimenez et al. 2005) with an integration step of 10^-4^ s.

**Figure 2.**
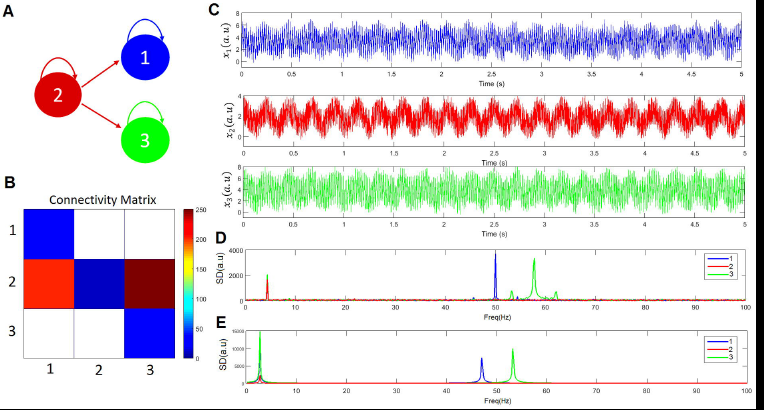
Three population toy model. A) The model comprises three neuronal populations labelled as ‘1’, ‘2’, and ‘3’, coloured in blue, red and green, respectively. This color legend is used across all panels in the figure. B) Connectivity matrix. C) Temporal dynamics of the three neuronal populations. D) Spectral density. E) Spectral density when substituting the sigmoid function with the linear function *S(x)* = *x*.

Figure 2C shows the temporal dynamics of the three populations and Figure 2D displays the corresponding spectral density. Population 2 oscillates at 4.40 Hz (theta band), whereas populations 1 and 3 have peaks at 50 and 57.8 Hz, respectively (gamma band). Because of the connections 2→1 and 2→3, there are peaks at 4.40 Hz in populations 1 and 3, and more importantly, there are secondary peaks at frequencies 50*Hz* ± 4.40 *Hz* and 57.8 *Hz* ± 4.40 *Hz* on both sides of these main peaks. This shows that the low frequency (4.40 Hz) is modulating the higher frequencies (50 and 57.8 Hz) and that there is theta-gamma PAC. According to the connections shown in Figure 2A, phases in populations 1 and 3 cannot modulate the amplitudes in populations 3 and 1, respectively. Thus, an appropriate method to study the generation of PAC should not detect any modulation between populations 1 and 3. We found that when the sigmoid function is replaced by the linear function *S(x)=x*, no modulation is obtained (Figure 2E).

Figure 3A shows the PAC computed using the four measures presented in section 2.2. Non-significant values are plotted in white. The four methods correctly detect that there is no PAC involving amplitudes in the gamma band in population 2 (there is no significant spectral peak at the gamma band, only white noise). However, according to ESC and Midx, there is significant PAC between the phases of the theta band in neuronal population 1 and the amplitudes of the gamma band in neuronal population 3, as well as PAC between the phases of the theta band in neuronal population 3 and the amplitudes of the gamma band in neuronal population 1. These results are expected because the signals in populations 1 and 3 are correlated, despite the fact that there is no connection between these populations. Regardless, cMI and cTE distinguished the correct effective interactions between the three populations.

**Figure 3.**
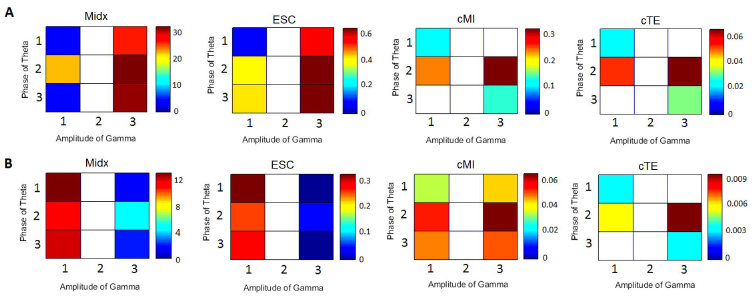
Causality vs correlation measures of PAC. A) Midx, ESC, cMI and TE. B) Midx, ESC, cMI and cTE when the noise is increased (*σ*_1_ = *σ*_2_ = *σ*_3_ = 10).

There are cases where cMI fails to estimate the correct connections. For instance, Figure 3B shows the results of increasing the noise (*σ*_1_ = *σ*_2_ = *σ*_3_ = 10), which caused the measures ESC, Midx and cMI to yield similar results and estimate a significant effective connection between populations 1 and 3 that did not exist. Regardless, cTE was still able to distinguish the correct pattern of connections despite the increase in the noise level. When we further increased the noise (*σ*_1_ = *σ*_2_ = *σ*_3_ = 30), no significant results were obtained for any of the four measures (not shown in the figure).

### 3.2 Generation of multiple PAC combinations

In this section, we study the generation of PAC in the cortical column circuit depicted in Figure 1. Since we are interested in the interaction between the rhythms produced by the nonlinear dynamics of the neuronal populations (not their correlation) and in the directionality of that interaction (from phases to amplitudes), we only compute cTE.

The values of the parameters used are shown in tables 2 and 3. White noise with a mean of zero and standard deviation *σ* = 1 was added to the external inputs. Five seconds of data were simulated and the first two seconds were discarded to avoid transient behavior. Thus, subsequent steps were carried out with the remaining three seconds.

**Table 3.**
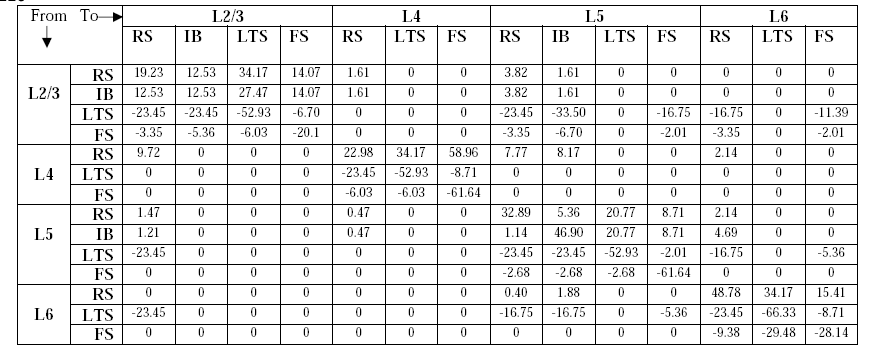
Standard values of the anatomically constrained effective connectivity matrix Γ*_nm_* (Figure 1B). All values represent anatomical (direct) connections. Nonzero values were manually tuned to produce peaks in the spectrum of *x_m_(t)* in all frequencies of interest (from delta to gamma) as well as an average LFP spectrum (Figure 6) consistent with experimental results (Maier, Adams et al. 2010, Buffalo, Fries et al. 2011). Values that are zero were taken from the literature (Neymotin, Jacobs et al. 2011).

| From<br>↓ | To→ | L2/3 |  |  |  | L4 |  |  | L5 |  |  |  | L6 |  |  |
| --- | --- | --- | --- | --- | --- | --- | --- | --- | --- | --- | --- | --- | --- | --- | --- |
|  |  | RS | IB | LTS | FS | RS | LTS | FS | RS | IB | LTS | FS | RS | LTS | FS |
| **L2/3** | **RS** | 19.23 | 12.53 | 34.17 | 14.07 | 1.61 | 0 | 0 | 3.82 | 1.61 | 0 | 0 | 0 | 0 | 0 |
|  | **IB** | 12.53 | 12.53 | 27.47 | 14.07 | 1.61 | 0 | 0 | 3.82 | 1.61 | 0 | 0 | 0 | 0 | 0 |
|  | **LTS** | -23.45 | -23.45 | -52.93 | -6.70 | 0 | 0 | 0 | -23.45 | -33.50 | 0 | -16.75 | -16.75 | 0 | -11.39 |
|  | **FS** | -3.35 | -5.36 | -6.03 | -20.1 | 0 | 0 | 0 | -3.35 | -6.70 | 0 | -2.01 | -3.35 | 0 | -2.01 |
| **L4** | **RS** | 9.72 | 0 | 0 | 0 | 22.98 | 34.17 | 58.96 | 7.77 | 8.17 | 0 | 0 | 2.14 | 0 | 0 |
|  | **LTS** | 0 | 0 | 0 | 0 | -23.45 | -52.93 | -8.71 | 0 | 0 | 0 | 0 | 0 | 0 | 0 |
|  | **FS** | 0 | 0 | 0 | 0 | -6.03 | -6.03 | -61.64 | 0 | 0 | 0 | 0 | 0 | 0 | 0 |
| **L5** | **RS** | 1.47 | 0 | 0 | 0 | 0.47 | 0 | 0 | 32.89 | 5.36 | 20.77 | 8.71 | 2.14 | 0 | 0 |
|  | **IB** | 1.21 | 0 | 0 | 0 | 0.47 | 0 | 0 | 1.14 | 46.90 | 20.77 | 8.71 | 4.69 | 0 | 0 |
|  | **LTS** | -23.45 | 0 | 0 | 0 | 0 | 0 | 0 | -23.45 | -23.45 | -52.93 | -2.01 | -16.75 | 0 | -5.36 |
|  | **FS** | 0 | 0 | 0 | 0 | 0 | 0 | 0 | -2.68 | -2.68 | -2.68 | -61.64 | 0 | 0 | 0 |
| **L6** | **RS** | 0 | 0 | 0 | 0 | 0 | 0 | 0 | 0.40 | 1.88 | 0 | 0 | 48.78 | 34.17 | 15.41 |
|  | **LTS** | -23.45 | 0 | 0 | 0 | 0 | 0 | 0 | -16.75 | -16.75 | 0 | -5.36 | -23.45 | -66.33 | -8.71 |
|  | **FS** | 0 | 0 | 0 | 0 | 0 | 0 | 0 | 0 | 0 | 0 | 0 | -9.38 | -29.48 | -28.14 |

Figure 4 presents the temporal evolution of the average PSP in each neuronal population. Time series coloured in red correspond to excitatory populations (L2RS, L2IB, L4RS, L5RS, L5IB, L6RS), whereas inhibitory populations (L2LTS, L4LTS, L5LTS, L6LTS) are represented in green. As seen in the figure, the generated signals show the characteristic ‘waxing and waning’ (i.e, amplitude modulation) observed in real EEG signals.

**Figure 4.**
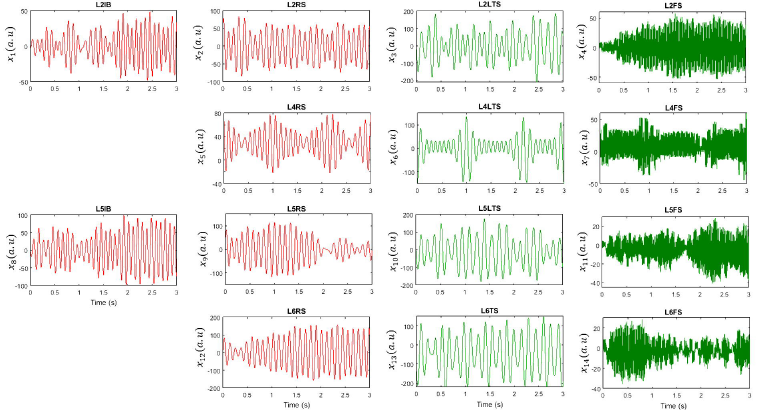
Simulated temporal evolution of the postsynaptic potentials for all populations for one realization of the model. Excitatory populations are depicted in red and inhibitory ones in green.

Figure 5 presents the normalized spectrum of the signals displayed in Figure 4. Excitatory populations are depicted in red and inhibitory populations are depicted in green. The six excitatory populations have their main spectrum peak in the alpha band, but they also present energy in the delta and theta bands. Slow inhibitory populations have the highest peak in the theta band, but also have energy in delta, alpha bands. Fast inhibitory populations were set to yield a peak in the gamma band, but due to the interaction with other populations they yield significant peaks in other frequencies as well, especially in the theta and alpha bands. This is evident when compared to the spectrum (in black) of the population when interactions between different populations are set to zero (Γ*_nm_* = 0, *n* ≠ *m*). Peaks in black correspond to the natural frequency of oscillation for the populations L2RS (9.00 Hz), L2IB (10.67 Hz), L2LTS (7.00 Hz), L2FS (58.67 Hz), L4RS (9.33 Hz), L4LTS (7.00 Hz), L4FS (87.00 Hz), L5RS (8.67 Hz), L5IB (9.67 Hz), L5LTS (7.00 Hz), L5FS (63.99 Hz), L6RS (7.67 Hz), L6LTS (7.33 Hz), and L6FS (59.67 Hz).

**Figure 5.**
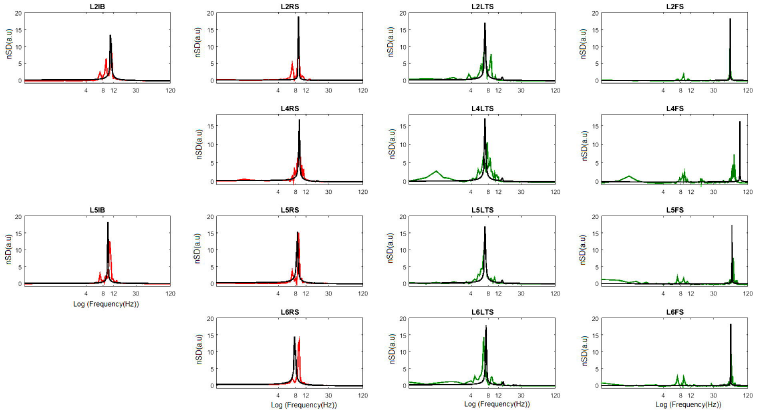
Normalized spectral density (nSD) of the postsynaptic potentials shown in Figure 3 obtained by subtracting the mean of the spectral density vector and dividing by the standard deviation. Excitatory populations are depicted in red and inhibitory ones in green. nSD coloured in black show the results when the connections between populations are set to zero.

To compare our simulated data with actual local field potential (LFP) data, we computed an approximation of the average LFP as the difference between the sum of excitatory and inhibitory activities in each layer.

Figure 6A displays the temporal dynamics of the LFP in each cortical layer and Figure 6B shows the corresponding spectral density. Thus, parameter values presented in tables 2 and 3 result in low frequency oscillations (delta, theta and alpha) with highest power in layers 5 and 6 while gamma oscillations have its main power in layer 2/3. This is in agreement with recent findings suggesting that gamma activity is predominant in superficial layers while lower frequencies are predominant in deep layers (Maier, Adams et al. 2010, Buffalo, Fries et al. 2011).

**Figure 6.**
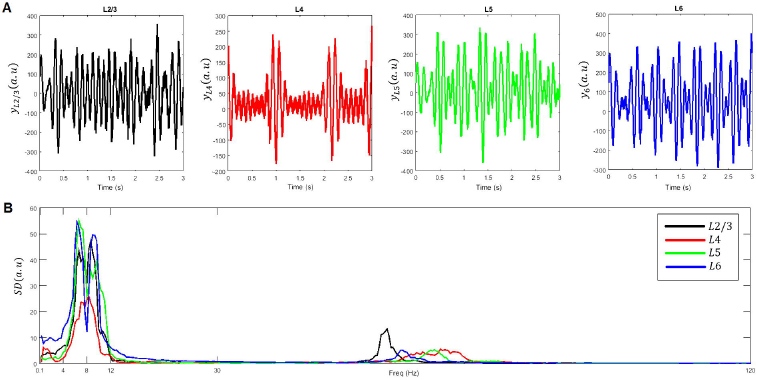
Laminar distribution of average LFP. A) Temporal dynamics in layers 2/3 (L2/3), 4(L4), 5(L5) and 6(L6). B) Spectral density (SD).

To test the existence of PAC, we filtered each time series in Figure 4 into five frequency bands from delta to gamma (see section 2.2) and applied the Hilbert transform to obtain instantaneous phases and amplitudes for each frequency band and each neuronal population.

Ten different PAC combinations between a low-frequency phase and a higher-frequency amplitude were computed using the cTE measure: delta-theta, delta-alpha, delta-beta, delta-gamma, theta-alpha, theta-beta, theta-gamma, alpha-beta, alpha-gamma, and beta-gamma. Each PAC combination consisted of a matrix of 14x14 cTE values representing all possible interactions between the 14 neuronal populations. To test the significance of these values, surrogate data was computed, followed by a multiple comparison analysis as described in section 2.2.

Results include nine out of the ten PAC combinations (Figure 7). The delta-beta PAC combination was not included since no significant values were obtained for the set of parameters used.

**Figure 7.**
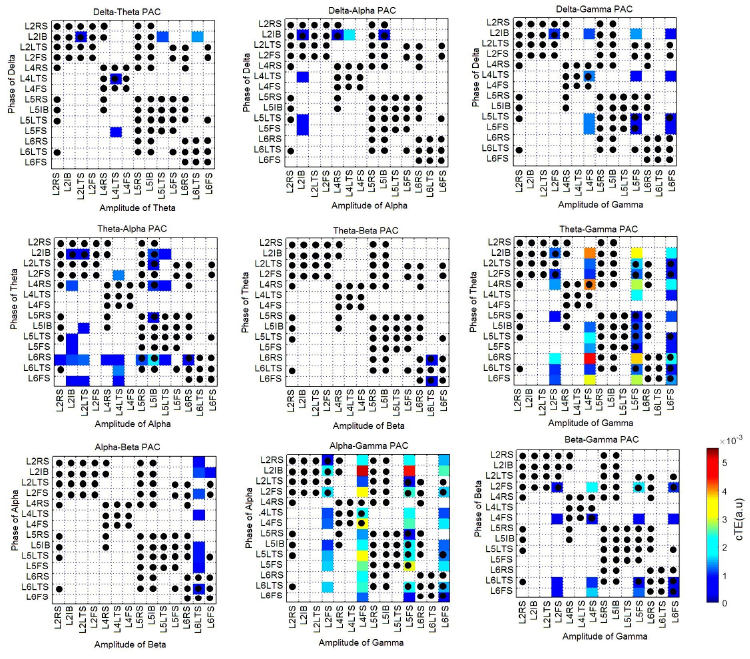
Phase-amplitude coupling (PAC) corresponding to the simulation presented in Figure 4. Non-significant values were set to zero and are depicted in white. Black dots indicate existing anatomical connections (see figure 1).

The strongest PAC value found was between the phase of the alpha band in L2IB and the amplitude of the gamma band in L4FS, which we will denote as L2IB→ L4FS. Other strong values found were theta-gamma (L6RS→L4FS, L4RS→L4FS, L2IB→L4FS, L6RS→L5FS), and alpha-gamma (L2IB→L5FS, L4FS→L4FS, L2FS→L4FS). Some of these values do not represent direct connections between the populations. For example, the strongest causal connection (L2IB→ L4FS) does not correspond to an anatomical (direct) connection (see Figure 1). Thus, we emphasize that PAC matrices (Figure 7) represent effective connections, which can correspond or not correspond with anatomical connections. To make this clearer, anatomical connections in Figure 7 are represented with black dots.

### 3.3 Parameter sensitivity analysis

Some of the parameters presented in table 2 were taken from the neural mass literature (Jansen and Rit 1995, Wendling, Bellanger et al. 2000), and parameters with no equivalent in the literature were assigned physiologically reasonable values. Thus, it is necessary to explore how changes in these parameters can affect PAC values. In this section, for the sake of simplicity, we focus on three PAC combinations which involve the gamma rhythm and have been of great interest in the literature: delta-gamma, theta-gamma, and alpha-gamma.

#### 3.3.1 Controlling the strength of PAC

We selected nine different parameters and explored how their change affected the strength of the PAC phenomenon. For each parameter we considered 100 different values and thus performed 100 different simulations. The parameters were: 1) a multiplying factor *η* = 0.3:0.3:3 controlling the global strength of the connectivity matrix (Γ_*nm*_ = *η*Γ*_nm_*), 2) the reciprocal of the time constant of RS populations (*k_RS_* = 5: 5: 500s^-1^), 3) the reciprocal of the time constant of IB populations (*k_IB_* = 5: 5: 500s^-1^), 4) the reciprocal of the time constant of LTS populations (*k_LTs_* = 5: 5:500s^-1^), 5) the reciprocal of the time constant of FS populations (*k_FS_* = 5: 5: 500s^-1^), 6) the external input to the L4RS population (*p*_5_ = 10:10:1000), 7) the external input to the L4FS population (*p*_7_ = 10:10:1000), 8) the gains of the six excitatory populations (*G*_E_ = *G*_1_ = *G*_2_ = *G*_5_ = *G*_8_ = *G*_9_ = *G*_12_ = 0.2:0.2: 20), and 9) the gains of the eight inhibitory populations (*G*_1_, = *G*_3_ = *G*_4_ = *G*_6_ = *G*_7_ = *G*_10_ = *G*_11_ = *G*_13_ = *G*_14_ = 0.3:0.3: 30).

Then, for each PAC combination we obtained 14x14x100= 19600 cTE values (although many of them are zero). We summarized that information by taking the strongest value found in each simulation, which results in a series of 100 values for each PAC combination. Figure 8A displays the mean and standard deviation of the 100-point series of the strongest PAC values for the three PAC combinations considered. In the figure, Delta-gamma PAC is depicted in orange, theta-gamma PAC in green, and alpha-gamma PAC in blue. Our results shows that for the three PAC combinations, the highest increase in cTE was obtained when changing the reciprocal time constants of LTS populations.

**Figure 8.**
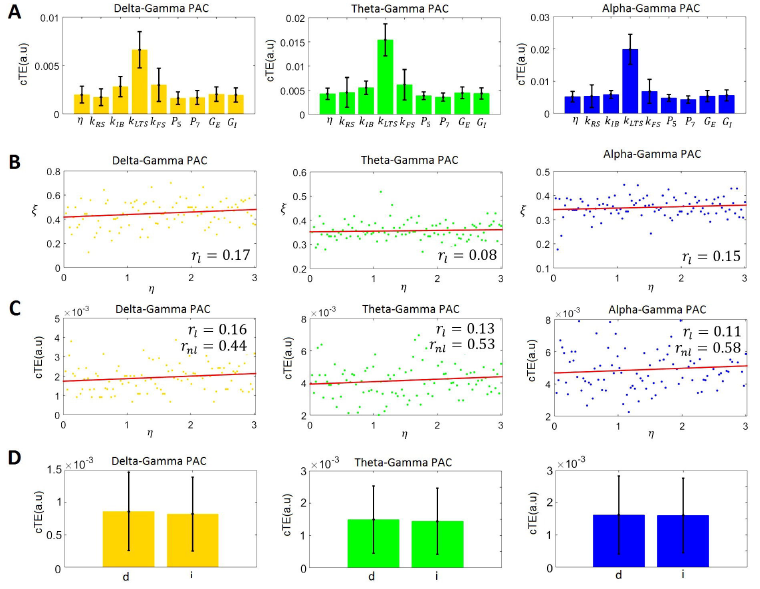
Exploring the parameter space for three different PAC combinations. A) Average cTE values for delta-gamma (orange), theta-gamma (green), and alpha-gamma (blue) PAC when considered 100 different values for nine different parameters: 1) a multiplying factor *η* = 0.3:0.3:3 controlling the global strength of the connectivity matrix (Γ*_nm_* = *η*Γ*_nm_*), 2) the reciprocal of the time constant of RS populations (*k_RS_* = 5: 5: 500*s*^-1^), 3) the reciprocal of the time constant of IB populations (*k_IB_* = 5: 5: 500*s*^-1^), 4) the reciprocal of the time constant of LTS populations (*k_LTS_* = 5: 5: 500*s*^-1^), 5) the reciprocal of the time constant of FS populations (*k_FS_* = 5: 5: 500*s*^-1^), 6) the external input to the L4RS population (*p*_5_ = 10:10:1000), 7) the external input to the L4FS population (*p*_7_ = 10:10:1000), 8) the gains of the six excitatory populations (*G_E_* ≡ *G*_1_ = *G*_2_ = *G*_5_ = *G*_8_ = *G*_9_ = *G*_12_ = 0.2:0.2:20), and 9) the gains of the eight inhibitory populations (*G*_7_ = *G*_3_ = *G*_4_ = *G*_6_ = *G*_7_ = *G*_10_ = *G*_1±_ = *G*_13_ = *G*_14_ = 0.3:0.3: 30). B) Plot of the fraction (*ξ*) of PAC connections that corresponded to anatomical connections versus *η*. C) Plot of cTE versus *η*. D) Average cTE values for direct and indirect PAC connections. Labels ‘d’, and ‘i’ correspond to direct and indirect PAC connections

The exploration of the parameter space is important because PAC has been suggested to be the carrier mechanism for the interaction of local and global processes in the brain, and is thus directly related to the integration of distributed information in the brain (Canolty and Knight 2010). Neuronal circuits can thus control the amount of information transmitted in the PAC phenomenon by changing the values of physiological parameters of specific populations.

#### 3.3.2 On the influence of the connectivity matrix Γ*_nm_* on PAC strength

An important problem in neuroscience is the link between structural and functional brain networks (Honey, Sporns et al. 2009, Stam, van Straaten et al. 2015). In the context of this work, it is of interest to study the influence of the connectivity matrix Γ*_nm_* on the generated PAC phenomenon.

For each of the 100 simulations we also computed the fraction (*ξ*) of PAC connections that corresponded to anatomical connections. The obtained ratios for delta-gamma, theta-gamma, and alpha-gamma, were 0.45 ± 0.12, 0.36 ± 0.04, and 0.35 ± 0.04, respectively. Figure 8B shows the plot of the ratios *ξ* versus the factor *η*, and the corresponding linear model fit. The linear correlation values (*r_l_*) between *ξ* and *η* for delta-gamma, theta-gamma, and alpha-gamma PAC were 0.17, 0.08, and 0.15, respectively. The values were significant with *α* = 0.05.

Figure 8C displays the series of 100 PAC values versus *η*. The solid line corresponds to the fit of a linear model. The linear correlation values between delta-gamma, theta-gamma, alpha-gamma and *η* were 0.16, 0.13, and 0.11, respectively. As expected, the linear correlation between the strength of the PAC phenomenon and the global strength of the connectivity matrix Γ*_nm_* was weak, although significant (with *α* = 0.05). We then performed a nonlinear regression analysis (equation 10) with *η* as the regressor and computed the nonlinear correlation coefficient (section 2.3). The nonlinear correlation values between delta-gamma, theta-gamma, alpha-gamma and *η* were 0.44, 0.53, and 0.58, respectively, showing that there is a strong nonlinear relationship between the strength of PAC and effective connectivity between the populations involved. The values were significant as tested with the surrogate data approach (section 2.2).

We also counted all significant PAC connections obtained in the 100 simulations. The vectors of significant connections for delta-gamma, theta-gamma, and alpha-gamma PAC comprised 595, 1464, and 1499, cTE values, respectively, for the PAC interactions that have a corresponding anatomical connection (direct interactions), and 740, 2656, and 2788 for the interactions without an anatomical equivalent (indirect interactions). The mean and standard deviations of these connections are presented in Figure 8D. Our results showed that for the three PAC combinations, the average strength of direct PAC interactions was slightly higher than indirect PAC interactions, but this difference was not statistically significant (as tested with a t-test, p<0.05).

Finally, we computed three local topological measures (section 2.4) for the network of 14 coupled neuronal populations (Fig 1A): *C_m_, E_m_*, and *B_m_*. The edges of the network were the absolute values of the connections between the populations (Figure 1B). We found that on average, indirect PAC interactions are made by populations with higher *C_m_* (Figure 9A), *E_m_* (Figure 9B), and *B_m_* (Figure 9C) than populations making direct connections, and populations not involved in PAC connections. Populations receiving indirect PAC connections had also on average higher topological measures than populations receiving direct interactions.

**Figure 9.**
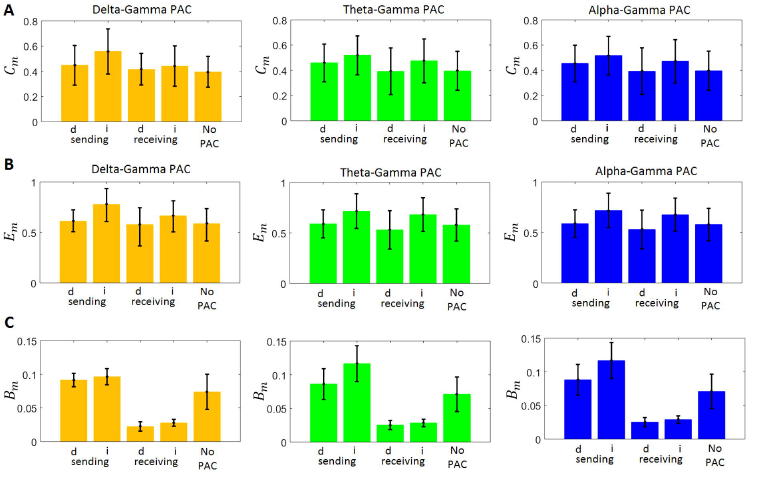
The link between local topological measures and PAC. A) local clustering coefficient (*C_m_*), B) local efficiency (*E_m_*), C) local betweenness centrality (*B_m_*). In all panels, labels ‘d’, and ‘i’ correspond to direct and indirect PAC connections, respectively. Populations can send and/or receive PAC interactions, or they can be not involved in the generation of PAC.

### 3.4 Generation of other types of CFC

The neural mass model presented in this paper can generate rich temporal dynamics. Studies of the dynamics generated by the Jansen and Rit model, which is the basis for our model, can be found elsewhere (Grimbert and Faugeras 2006, Faugeras, Veltz et al. 2009, Spiegler, Kiebel et al. 2010). In this paper we focused on PAC, but this is only one type of the general phenomenon of CFC which results from nonlinearities in brain dynamics. It is thus not unexpected to find other types of CFC in the signals generated by our model (for example, the temporal dynamics of L5RS in Figure 4 correspond to frequency modulation). In addition to PAC, other types of CFC such as AAC, PPC, and phase-frequency coupling (PFC) have been explored in the literature (Jirsa and Muller 2013, Hyafil, Giraud et al. 2015) and could all be calculated using equation (8) after replacing *a_nl_(t)* and *φ_mk_(t)* with the appropriate time series.

Recently, the analysis of resting state electrocorticography (ECoG) data revealed that the amplitude of gamma oscillations can drive the phase of alpha oscillations, i.e, APC (Jiang, Bahramisharif et al. 2015). Although this experimental result may seem surprising, it is consistent with theoretical results in the NMM literature. Specifically, starting with a network of Wilson and Cowan oscillators, equations for the instantaneous phases were obtained which depended on the instantaneous amplitudes of the oscillators in the network (Daffertshofer and van Wijk 2011). Thus, by setting different natural frequencies for the oscillators in the network, it is possible to obtain not only PAC but other types of CFC. To test the existence of APC we computed:

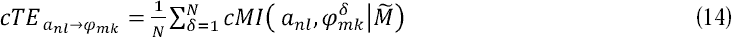
where 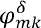 is derived from the phase time series *φ_mk_* at *δ* steps into the future, i.e. 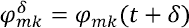, and 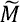 is a matrix comprising all phases and amplitudes in all populations, except *a_ni_*. Figure 10 shows the APC estimated from the simulated data shown in Figures 4 and 5, with the strongest values corresponding to the gamma-delta and gamma-theta APC combinations. Our simulation supports the existence of APC as recently proposed (Jiang, Bahramisharif et al. 2015).

**Figure 10.**
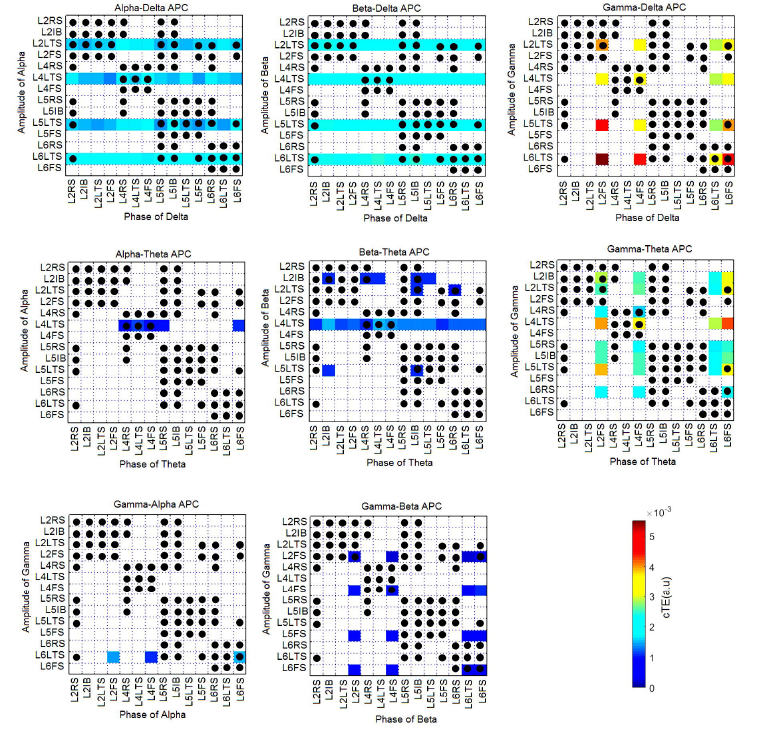
Amplitude-phase coupling (APC) corresponding to the simulation presented in Figures 4 and 5. Non-significant values were set to zero and are depicted in white. Black dots indicate existing anatomical connections (see figure 1).

### 3.5 Mechanisms mediating indirect PAC interactions

Due to the prevalence of indirect PAC interactions (Figure 7) it is necessary to investigate the mechanisms generating them. As mentioned in section 3.4, theoretical results have shown that amplitudes and phases are coupled even under weak coupling (Guckenheimer and Holmes 1997, Daffertshofer and van Wijk 2011). Thus, we expect that indirect information transfer from the phase of one population to the amplitude of the receiving population is mediated by direct interactions between phases and amplitudes via the connectivity matrix Γ_*nm*_. It should be noted that although PAC typically refers to the interaction between the phase of a low frequency rhythm and the amplitude of a higher frequency rhythm, interactions between phases and amplitudes of the same frequency or between the phase of a high frequency and the amplitude of a low frequency are also possible.

In this section, to explore the mechanism mediating indirect PAC interactions, we focused on three populations labelled from 1 to 3 (Figures 11 and 12) and connected in such a way that there was not a direct connection between populations 1 and 3:1→2→3. Population 1 oscillated in the theta band and population 3 oscillated in the gamma band. Two different cases were considered for population 2. Case I (Figure 11): Population 2 oscillated with a frequency lower (delta band) than population 1, and Case II (Figure 12): population 2 oscillated with a frequency higher (beta band) than population 1. Five different types of CFC were computed between the three populations (PAC, APC, PPC, AAC, and PFC) while varying the connectivity parameters between populations 1 and 2 (*c*_12_) and between populations 2 and 3 (*c*_23_). SFC was also considered and labelled in the same way as the CFC interactions. For example, *PPC*_*θ*_1_*θ*_2__, is the cTE from the phase of theta in population 1 to the phase of theta in population 2. To test the significance of these values, surrogate data was computed, followed by a multiple comparison analysis (section 2.2). As a control, we computed the cTE from populations 2 and 3 to population 1 for all possible types of interactions (such as *PPC*_*θ*_2_*θ*_1__), and confirmed they were not statistically significant.

**Figure 11.**
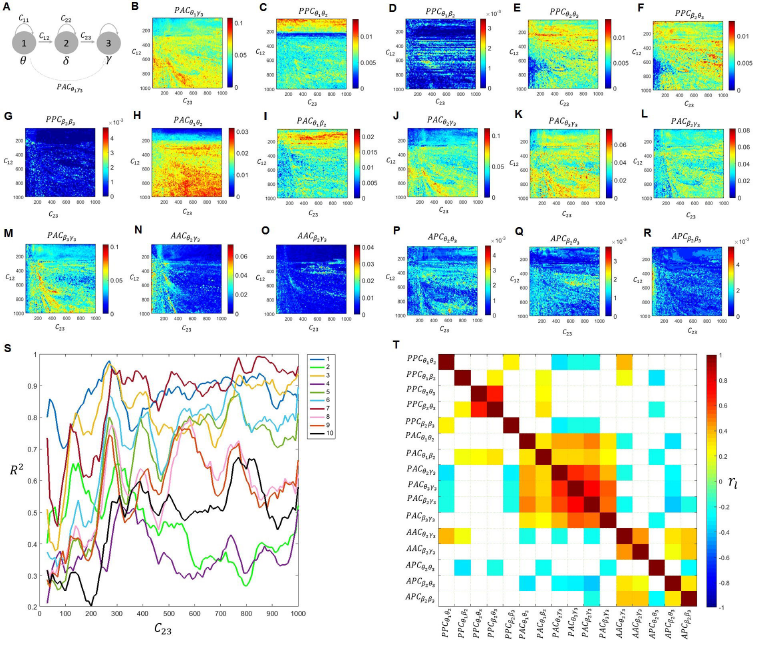
Mechanisms mediating indirect PAC connections (Case I). A) Three population toy model comprising three neuronal populations labelled as ‘1’, ‘2’, and ‘3’, oscillating in the theta (*θ*), delta (*δ*), and gamma (*γ*) bands. B) PAC involving the phase of theta in population 1 and the amplitude of gamma in population 3 (*PAC*_*θ*_1_*γ*_3__) obtained when varying the connectivity parameters between populations 1 and 2 (*c*_12_ = 30:4:1000) and between populations 2 and 3 (*c*_23_ = 30:4:1000). Panels C to R, displays the 16 predictors used in the ten models explored (table 4). S) Coefficient of determination (R^2^) for the ten models explored (table 4). T) Correlation coefficient between the 16 predictors.

**Figure 12.**
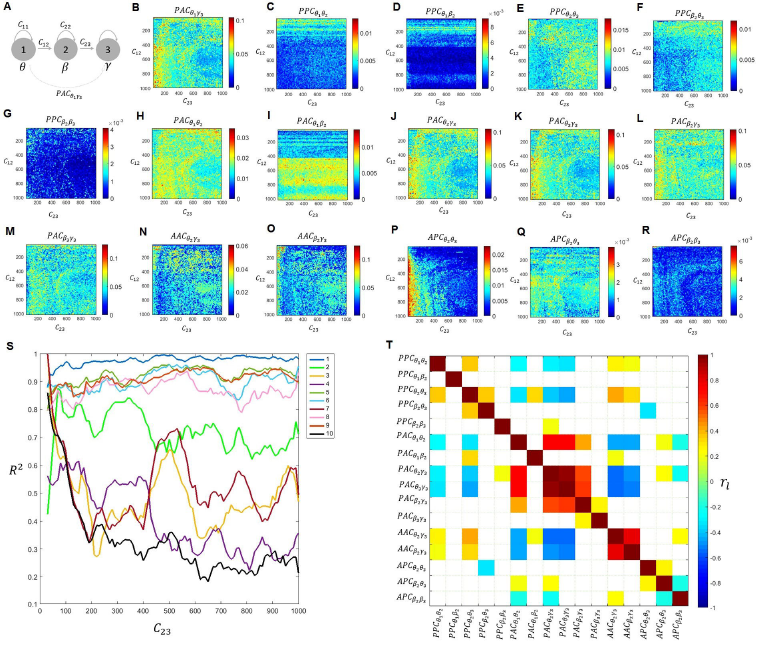
Mechanisms mediating indirect PAC connections (Case II). A) Three population toy model comprising three neuronal populations labelled as ‘1’, ‘2’, and ‘3’, oscillating in the theta (*θ*), beta (*δ*), and gamma (*γ*) bands. B) PAC involving the phase of theta in population 1 and the amplitude of gamma in population 3 (*PAC*_*θ*_1_*γ*_3__) obtained when varying the connectivity parameters between populations 1 and 2 (*c*_12_ = 30:4:1000) and between populations 2 and 3 (*c*_23_ = 30:4:1000). Panels C to R, display the 16 predictors used in the ten models explored (table 4). S) Coefficient of determination (R^2^) for the ten models explored (table 4). T) Correlation coefficient between the 16 predictors.

**Table 4.** Indirect PAC modeled as a cascade of direct CFC and SFC in a three population network. Two cases were considered: population 2 oscillates in the beta (*β*) band (Case I), and population 2 oscillates in the delta (*δ*) band (Case II). Populations 1 and 3 always oscillate in the theta (*θ*) and gamma (*γ*) bands, respectively.

| Model | Case I | Case II |
| --- | --- | --- |
| 1 | $PAC_{\theta_1\gamma_3} = PPC_{\theta_1\theta_2} + PAC_{\theta_2\gamma_3}$ | $PAC_{\theta_1\gamma_3} = PPC_{\theta_1\theta_2} + PAC_{\theta_2\gamma_3}$ |
| 2 | $PAC_{\theta_1\gamma_3} = PAC_{\theta_1\theta_2} + AAC_{\theta_2\gamma_3}$ | $PAC_{\theta_1\gamma_3} = PAC_{\theta_1\theta_2} + AAC_{\theta_2\gamma_3}$ |
| 3 | $PAC_{\theta_1\gamma_3} = PPC_{\theta_1\beta_2} + PAC_{\beta_2\gamma_3}$ | $PAC_{\theta_1\gamma_3} = PPC_{\theta_1\delta_2} + PAC_{\delta_2\gamma_3}$ |
| 4 | $PAC_{\theta_1\gamma_3} = PAC_{\theta_1\beta_2} + AAC_{\beta_2\gamma_3}$ | $PAC_{\theta_1\gamma_3} = PAC_{\theta_1\delta_2} + AAC_{\delta_2\gamma_3}$ |
| 5 | $PAC_{\theta_1\gamma_3} = PPC_{\theta_1\theta_2} + PPC_{\theta_2\theta_3} + PAC_{\theta_3\gamma_3}$ | $PAC_{\theta_1\gamma_3} = PPC_{\theta_1\theta_2} + PPC_{\theta_2\theta_3} + PAC_{\theta_3\gamma_3}$ |
| 6 | $PAC_{\theta_1\gamma_3} = PPC_{\theta_1\beta_2} + PPC_{\beta_2\theta_3} + PAC_{\theta_3\gamma_3}$ | $PAC_{\theta_1\gamma_3} = PPC_{\theta_1\delta_2} + PPC_{\delta_2\theta_3} + PAC_{\theta_3\gamma_3}$ |
| 7 | $PAC_{\theta_1\gamma_3} = PPC_{\theta_1\beta_2} + PPC_{\beta_2\beta_3} + PAC_{\beta_3\gamma_3}$ | $PAC_{\theta_1\gamma_3} = PPC_{\theta_1\delta_2} + APC_{\delta_2\delta_3} + PAC_{\delta_3\gamma_3}$ |
| 8 | $PAC_{\theta_1\gamma_3} = PAC_{\theta_1\theta_2} + APC_{\theta_2\theta_3} + PAC_{\theta_3\gamma_3}$ | $PAC_{\theta_1\gamma_3} = PAC_{\theta_1\theta_2} + APC_{\theta_2\theta_3} + PAC_{\theta_3\gamma_3}$ |
| 9 | $PAC_{\theta_1\gamma_3} = PAC_{\theta_1\beta_2} + APC_{\beta_2\theta_3} + PAC_{\theta_3\gamma_3}$ | $PAC_{\theta_1\gamma_3} = PAC_{\theta_1\delta_2} + APC_{\delta_2\theta_3} + PAC_{\theta_3\gamma_3}$ |
| 10 | $PAC_{\theta_1\gamma_3} = PAC_{\theta_1\beta_2} + APC_{\beta_2\beta_3} + PAC_{\beta_3\gamma_3}$ | $PAC_{\theta_1\gamma_3} = PAC_{\theta_1\delta_2} + APC_{\delta_2\delta_3} + PAC_{\delta_3\gamma_3}$ |

There are several pathways that could transfer information from the phase of the theta oscillation in population 1 to the amplitude of the gamma oscillation in population 3 (*PAC*_*θ*_1_*γ*_3__). For example, a simple model could involve *PAC*_*θ*_1_*θ*_2__ followed by PAC between the theta rhythm in population 2 and the gamma rhythm in population 3: *PAC*_*θ*_1_*γ*_3__ = *PAC*_*θ*_1_*θ*_2__ + *PAC*_*θ*_2_*γ*_3__. A more complicated one is: *PAC*_*θ*_1_*γ*_3__ = *PAC*_*θ*_1_*δ*_3__ + *PAC*_*δ*_2_*δ*_3__ + *PAC*_*δ*_3_*γ*_3__.

To compare different models (Table 4), we fitted a linear regression to *PAC*_*θ*_1_*γ*_3__ and computed the coefficient of determination (*R^2^*) as a function of parameter *c*_23_. For Case I we found that the three best models were Model 1 (*PAC*_*θ*_1_*γ*_3__ = *PAC*_*θ*_1_*θ*_2__ + *PAC*_*θ*_2_*γ*_3__), Model 7 (*PAC*_*θ*_1_*γ*_3__ = *PAC*_*θ*_1_*δ*_2__ + *PAC*_*δ*_2_*δ*_3__ + *PAC*_*δ*_3_*γ*_3__), and Model 3 (*PAC*_*θ*_1_*γ*_3__ = *PAC*_*θ*_1_*δ*_2__ + *PAC*_*δ*_2_*γ*_3__), with Model 1 being the best model for low values of *c*_23_, and Model 7 for the high values. On the other hand, we obtained the opposite behavior for Case II, i.e, Model 7 was the best model for low *c*_23_ values, whereas Model 1 was better at explaining *PAC*_*θ*_1_*γ*_3__ for high *c*_23_ values. The correlation between the 16 CFC and SFC variables involved in the ten models is displayed in the last panel for both cases (Figures 11 and 12). Although we found significant PFC combinations, models involving these combinations were very weak predictors of *PAC*_*θ*_1_*γ*_3__ (with R^2^ <0.08 in all cases) and were thus not included in Table 3 and Figures 11 and 12.

## 4. Discussion

We have proposed a neural mass model that captures the phase-amplitude coupling between layers in a cortical column. The model comprises fourteen interconnected neuronal populations distributed across four cortical layers (L2/3, L4, L5 and L6). We omitted layer 1, because it does not include somas (Binzegger, Douglas et al. 2004). Based on experimental reports on the strength of the inputs to each layer (Binzegger, Douglas et al. 2004, Jellema, Brunia et al. 2004), we only considered external inputs to the RS and FS populations in layer 4, thus neglecting possible external inputs to other layers.

According to our results, the parameters with the strongest influence on the strength of PAC were the time constants (especially the ones for LTS, FS, and IB populations). As expected, in order to generate PAC, nonlinearities in the model are essential. As was shown in Figure 2E, when the sigmoid function was substituted with a linear function, no modulation was obtained. Additionally, the strength of PAC was best modeled by a nonlinear regression of the connectivity values instead of a linear regression. Thus, our results show that the nonlinear interaction of neuronal populations (via the sigmoid function and the connectivity matrix) can produce PAC combinations with frequencies different from the natural frequencies of the oscillators involved. Our model of oscillators with natural frequencies in the theta, alpha and gamma bands was able to produce significant PAC involving delta and beta rhythms, including delta-alpha, delta-beta, delta-gamma, theta-beta, alpha-beta, and beta-gamma. Interestingly, some peaks in the beta band are harmonics of theta and alpha oscillations, such as the beta peak at 19 Hz in the spectrum of L4FS in Figure 5. Due to the interaction between the populations, there is a statistically significant PAC from the phase of beta in L4FS to the amplitude of gamma in L2FS, L4FS, L5FS and L6FS. Note that of these PAC interactions, only L4FS→L4FS corresponds to an anatomical connection (Figure 1B and table 3). If we take into account all PAC combinations in Figure 7, less than 40% of all significant PAC values (60/160=37.50%) corresponded to anatomical connections. This suggests that although effective connections are constrained by direct (anatomical) connections (Sotero, Bortel et al. 2010) additional factors are needed to fully explain the link between anatomical and effective connectivity. Interestingly, our numerical simulations showed that on average the strength of the PAC phenomenon mediated by direct and indirect connections is approximately the same. However, local topological measures such as clustering coefficient, efficiency, and betweenness centrality were the highest for populations making indirect connections when compared to populations making direct PAC connections, to populations receiving PAC connections, and to populations not involved on the generation of PAC. This suggest that the topology of cortical circuits shapes the generation of the PAC phenomenon. This is another factor to consider when studying the origin of PAC during neurodegenerative disorders known to affect both local and global brain circuitry.

One limitation of the present approach is that model parameters are loosely constrained from existing neurophysiological data. Thus, although our model provides insight about the emergence of PAC in a complex network whose spectral and connectivity properties resemble that of a cortical column, specific conclusions should await to more knowledge of these data.

### 4.1 Comparison with previous models of PAC generation

The first computational models of PAC generation were realistic models of the theta-gamma coupling in the hippocampus (Kopell, Boergers et al. 2010). These models considered networks of hundreds of interconnected neurons which were individually modeled by either a single compartment (White, Banks et al. 2000) or realistically represented by multiple compartments for the soma, axon, and dendrites (Tort, Rotstein et al. 2007). A practical disadvantage of this approach is that it needs high computational power, but more importantly, the use of such realistic models produces hundreds or thousands of variables and parameters, making it difficult to determine their influence on the generated average network characteristics. This is especially critical if we are interested in analyzing the link of PAC and mesoscopic phenomena like functional magnetic resonance signals (Wang, Saalmann et al. 2012). The analysis of multiple PAC combinations as done in this paper would be even more difficult with realistic networks. By comparison, our model of one cortical column comprised only 14 second-order (or 28 first-order) differential equations, which can be easily solved using any modern personal computer.

Additionally, previous models of PAC generation, both the ones based on realistic networks (Kopell, Boergers et al. 2010) and the ones based on neural mass models (Onslow, Jones et al. 2014) studied the phenomenon in a qualitative way, such that they did not actually compute a PAC measure but limited their analysis to the generation of temporal dynamics resembling PAC. This makes it difficult to compare their results with our quantitative approach based on information theory.

### 4.2 Indirect PAC connections can be predicted by a cascade of direct CFC and interactions within the same frequency band

As a unifying theory of EEG organization, it has been proposed that the EEG is hierarchically organized such that the delta phase modulates the theta amplitude, and the theta phase modulates the gamma amplitude (Lakatos, Shah et al. 2005). It was also proposed that this oscillatory hierarchy controls baseline excitability and that the hierarchical organization of ambient oscillatory activity allows the auditory cortex to structure its temporal activity pattern to optimize the processing of rhythmic inputs. Recent findings suggest a somewhat different hierarchy of oscillatory activity with regard to these frequency bands (Sotero, Bortel et al. 2015). Sotero et al. did not observe PAC between the delta and theta bands in rat area S1FL: PAC was statistically significant between the phases of the delta and theta bands and the amplitudes of the beta and gamma bands, but not between the phase of the delta band and the amplitude of the theta band. Their data support specific PAC interactions, but not a clear hierarchical PAC structure. The differences between Lakatos et al.’s findings and Sotero et al.’s findings are consistent with their proposal that the hierarchical structure found in the auditory cortex may support processing of rhythmic auditory stimuli, which are less common in natural somatosensory stimuli to the forepaw. Both studies were restricted to PAC and did not explore whether the oscillatory hierarchy might involve other types of CFC. While historically PAC and PPC have been the subject of most experimental and modeling studies, other types of CFC are attracting increasing interest (Jirsa and Muller 2013, Hyafil, Giraud et al. 2015, Jiang, Bahramisharif et al. 2015). Our results show that indirect PAC is better understood if analyzed along with direct PAC and other types of direct CFC and SFC connections. Our simulations do not suggest a specific oscillatory hierarchy like the one proposed by Lakatos et al., but multiple contributing cascades of CFC and SFC. Future analysis of experimental data will need to determine the functional importance of these different possible pathways.

### 4.3 cTE as a unified approach to estimate CFC

In this work, we used the average cTE, computed using the conditional mutual information (Palus, Komarek et al. 2001, Palus and Stefanovska 2003) to measure the influence of the phase of a low frequency rhythm on the amplitude of a higher frequency rhythm, and used it as an index of PAC. A known limitation of the cTE approach is that it requires long time series (Hlavackova-Schindler, Palus et al. 2007). For this reason, we used time series comprising of 30,000 time instants. Recent studies have shown that cTE is biased as its values depend on the autodependency coefficient in a dynamical system (Runge, Heitzig et al. 2012). Conditional TE was chosen over pairwise mutual information (MacKay 2003) or the pairwise information flow (Liang 2014) because pairwise analysis cannot distinguish between connectivity configurations such as [X→Y, X→Z, Z→Y] and [X→Z, Z→Y] (Ding 2006).

An advantage of measures based on information theory is that they are model-free. This is in contrast to other measures like Granger causality, which are based on autoregressive models (Seth, Barrett et al. 2015). Furthermore, Granger causality should not be applied to band-passed signals because the filtering process produces a large increase in the empirical model order, which often results in spurious results (Barnett and Seth 2011). Another advantage of the cTE measure is that it can be used to estimate any type of CFC, not just PAC. Thus, it provides a unified measure to study the CFC phenomenon.

cTE has often been given a causal interpretation, however a more recent point of view (Lizier and Prokopenko 2010) suggests that cTE should be interpreted as predictive information transfer, i.e. the amount of information that a source variable adds to the next state of the destination variable. Ultimately, interventions are required to detect causal interactions (Pearl 2000). This formalism is used in a causal measure called information flow (Ay and Polani 2008), which is also based on the cMI.

## Acknowledgements

The author thanks Lazaro Sanchez for his comments on a previous version of the manuscript, and Erica A. Baines for English editing. This work was partially supported by grant RGPIN-2015-05966 from Natural Sciences and Engineering Research Council of Canada.

